# Inheritance of associative memories in *C. elegans* nematodes

**DOI:** 10.1101/2020.09.08.287318

**Authors:** Noa Deshe, Yifat Eliezer, Lihi Hoch, Eyal Itskovits, Shachaf Ben-Ezra, Alon Zaslaver

## Abstract

The notion that associative memories may be transmitted across generations is intriguing, yet controversial. Here, we trained *C. elegans* nematodes to associate an odorant with stressful starvation conditions, and surprisingly, this associative memory was evident two generations down of the trained animals. The inherited memory endowed the progeny with a fitness advantage, as memory reactivation induced rapid protective stress responses that allowed the animals to prepare in advance for an impending adversity. Sperm, but not oocytes, transmitted the associative memory, and the inheritance required H3K9 and H3K36 methylations, the small RNA-binding Argonaute NRDE-3, and intact neuropeptide secretion. Remarkably, activation of a single chemosensory neuron sufficed to induce a serotonin-mediated systemic stress response in both the parental trained generation and in its progeny. These findings challenge long-held concepts by establishing that associative memories may indeed be transferred across generations.

## Introduction

The capacity to form memories allows individuals to make educated decisions based on past experiences. A key memory paradigm is known as the associative memory, where a link between two seemingly unrelated cues is established. A classic example is the Pavlovian dogs who associated food (the unconditioned stimulus, US) with the sound of a bell (conditioned stimulus, CS). As a result, these dogs started salivating in response to the mere sound of the bell in expectations of the associated food (Pavlov and Thompson, 1902).

Associative memories become particularly advantageous when the associated US is unfavorable (*e.g.,* a shock or a stress) as encountering the CS induces memory retrieval and individuals can consequently anticipate the impending adversity and respond to it in advance (Clayton et al., 2003; Eliezer et al., 2019). A compelling idea is that these valuable associative memories may be epigenetically transmitted to subsequent generations and provide them with a fitness advantage upon memory reactivation.

A rich body of literature describes how extreme life experiences modulate physiology and behavior of subsequent generations (Agrawal et al., 1999; Carone et al., 2010; Gapp et al., 2014; Öst et al., 2014; Rechavi et al., 2014; Bohacek and Mansuy, 2015; Jobson et al., 2015; Kishimoto et al., 2017). However, evidence for inheritance of associative memories is scarce: In rodents, mice trained to associate an odor with an aversive electric shock, transferred the acquired memory to their descendants (Dias and Ressler, 2014). These memory traces were evident even in the F3 generation, indicating of transgenerational, rather than intergenerational, inheritance mechanisms (Heard and Martienssen, 2014; Dias et al., 2015; Perez and Lehner, 2019). In *C. elegans* worms, repeated olfactory imprinting for at least four generations is then stably inherited through multiple successive generations (Remy, 2010). Furthermore, worms grown on the pathogenic bacteria pseudomonas learn to avoid bacterial-derived odorants, an information that is passed on to the next generation (Pereira et al., 2020). Interestingly, RNAs originating from the pathogenic bacteria induce avoidance behavior that persists four generations downstream of the parental generation that experienced the bacteria (Moore et al., 2019; Kaletsky et al., 2020).

Three main mechanisms may underlie transmittance of epigenetic information across generations: DNA methylation, chromatin modifications, and small RNAs (Heard and Martienssen, 2014; Liberman et al., 2019; Perez and Lehner, 2019). The latter two presumably work in concert where small RNAs are thought to direct histone-modifying enzymes to specific chromatin sites and by this modulate gene expression programs (Burton et al., 2011; Rechavi and Lev, 2017; Skvortsova et al., 2018; Weiser and Kim, 2019; Liberman et al., 2019; Duempelmann et al., 2020).

In that respect, *C. elegans* worms offer a powerful model to studying inheritance of associative memories: Their short 3-day life cycle enables rapid interrogation of multiple consecutive generations and their compatibility with a myriad of genetic manipulations already extracted evolutionarily conserved mechanisms that promote epigenetic inheritance (Burton et al., 2011; Rankin, 2015; Klosin et al., 2017; Rechavi and Lev, 2017; Minkina and Hunter, 2018; Weiser and Kim, 2019; Baugh and Day, 2020). Furthermore, equipped with a compact neural network consisting of 302 neurons (White et al., 1986; Cook et al., 2019), *C. elegans* worms form various types of associative memories (Ardiel and Rankin, 2010; Sasakura and Mori, 2013; Pritz et al., 2020), and the stereotypic animal-to-animal anatomy makes them an appealing system for mapping individual memory-storing neurons (Jin et al., 2016; Eliezer et al., 2019; Pritz et al., 2020).

Crucially, recent studies, primarily in *C. elegans* nematodes, demonstrated that the famous ‘Weismann barrier’ is breached and information from somatic neurons is transmitted to the germline to affect the physiology and behavior of subsequent generations (Remy, 2010; Gapp et al., 2014; Devanapally et al., 2015; Posner et al., 2019; Kaletsky et al., 2020; Pereira et al., 2020; Perez et al., 2020). These findings pave the way to further challenge current notions and ask if and how associative memories may be transmitted across generations.

Here we established that associative memories are indeed transferred to subsequent generations. We trained *C. elegans* worms to associate an attractive odorant with a stressful starvation, and remarkably, both the F1 and F2 descendants, initiated fast stress responses upon exposure to the odorant. This inheritance was mediated via the sperm, but not the oocytes, and depended on H3K9 and H3K36 methylations, the small RNA-binding Argonaute NRDE-3, as well as on intact neuropeptide secretion. Notably, the same chemosensory neuron that encoded the memory in the trained parental generation also stored the information in the descendants, and its sole activation sufficed to induce systemic stress responses. Finally, the inherited memory conferred the progeny a fitness advantage, thus providing the evolutionary basis for the emergence of this valuable capacity in animals.

## Results

In *C. elegans*, the mere reactivation of a stressful memory induces a systemic stress response (Eliezer et al., 2019). This is manifested by the rapid translocation of the general stress factor DAF-16/FOXO to nuclei of the gonad sheath cells. Anatomically, these cells are intimately associated with the germ cells (Greenstein, 2005), raising the possibility that memory information may be transmitted to the progeny.

To study this possibility, we trained hermaphroditic worms to form an associative memory by coupling the favorable odorant isoamyl alcohol (IAA) with starvation (**Figure 1A** and Methods). Indeed, odor-evoked memory reactivation induced a rapid translocation of the DAF-16/FOXO to cells’ nuclei, indicating that the trained P0 generation worms formed robust memory traces (**Figure 1B-C,E**). Surprisingly, F1 descendants of these trained P0 hermaphrodites inherited the aversive associative memory, as exposing them to IAA also induced a rapid translocation of the DAF-16/FOXO to cells’ nuclei (**Figure 1D-E and supplementary figures 1-2)**. Moreover, the stressful associative memory was also transmitted to the F2 generation, who similarly showed rapid DAF-16/FOXO translocation to cells’ nuclei upon exposure to the conditioned stimulus IAA.

**Figure 1.**
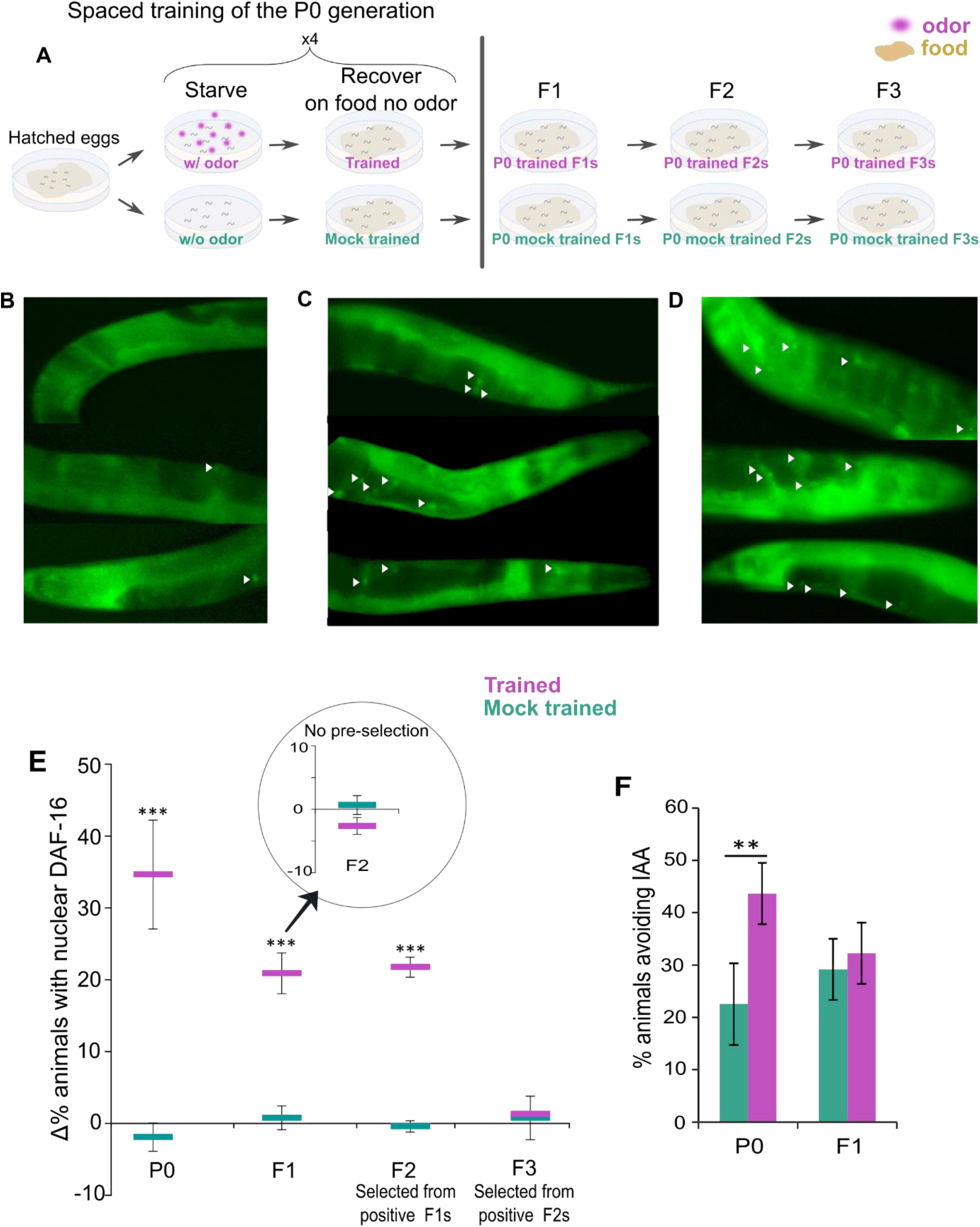
Aversive associative memories are transferred to the F1 and F2 generations. (**A**) A spaced-training paradigm was used to form the aversive associative memory: P0-generation worms were intermittently starved four times in the presence (trained) or the absence (mock trained) of the conditioned odorant IAA (see Methods). A full recovery on food in the absence of IAA followed each of the starvation steps. Subsequent generations F1, F2 and F3 were grown in normal satiety conditions. (**B-D**) Odor-evoked memory reactivation induced a rapid (<20 minutes) translocation of DAF-16/FOXO to cells’ nuclei. Shown are representative animals of: **(B)** Mock-trained P0 animals. **(C)** Trained P0 animals. **(D)** F1 progeny of P0 trained animals. Translocation of DAF-16/FOXO was observed primarily in the gonad sheath cells (white arrowheads) and was visualized using a translational fusion of the protein (Henderson and Johnson, 2001). We scored animals as initiating a stress response if at least six gonad sheath cells (in both gonad arms) showed nuclear DAF-16/FOXO localization. Mock-trained animals typically had <=1 cells with nuclear DAF-16/FOXO (note that the images show only one arm of the gonads; An extensive statistical validation for this scoring approach is found in Eliezer et al., figure 1 (Eliezer et al., 2019)). Each worm shown was imaged separately, cropped along its edges, and stacked one above the other. (**E**) The associative memory is transmitted to the F1 and the F2, but not to the F3, generations. Quantification of the stress response is based on the percent of animals detected with nuclear DAF-16/FOXO following odor-evoked memory reactivation (see supplementary figure 1 for the raw data). As transmittance efficiency diminished across generations, we scored F2s (F3s) originating from F1 (F2) generation worms that showed odor-evoked memory reactivation (noted as selected worms). The data in the inset is the % stressed worms in the F2 generation when picking progeny from the entire F1 population without preselecting memory-positive F1s. Each data point is the mean of at least three independent experimental repeats, each scoring ~50 animals. Error bars indicate SEM. ***p < 0.0005 (proportion test). (**F**) Trained P0 animals, but not their F1 descendants, withdrew when presented with the conditioned stimulus IAA. Forward-moving animals were presented with IAA, and were scored as avoiding IAA if they stopped and backed within 3 seconds. Shown are the means±SEM of 3–5 independent experimental repeats, each scoring ~50 animals. **p < 0.005 (proportion test).

The efficiency of the inheritance considerably diminished across generations (**Figure 1E and supplementary figure 3**), and to detect F2 animals that inherited the memory, we analyzed progeny of F1 animals that showed memory reactivation capacities (**Figure 1E**, compare to inset). We could not detect F3-generation worms that inherited the memory, even when pre-selecting progeny of memory-positive F2 worms (**Figure 1E**).

Next, we asked whether odor-evoked memory reactivation also triggers behavioral responses. For this, we trained animals as previously described (**Figure 1A**) and quantified their withdrawal following exposure to the conditioned stimulus IAA. Trained P0 worms withdrew upon exposure to IAA, however, their F1 descendants did not exhibit this avoidance behavior (consequently, F2 descendants were not assayed, **figure 1F**). Taken together, odor-evoked memory reactivation triggers both behavioral and physiological responses in the trained P0 generation, but only the physiological capacity to initiate a stress response upon memory retrieval was inherited by the F1 and the F2 generations.

Odor-evoked reactivation of stressful memories induces rapid protective stress responses that allow animals to prepare in advance for impending adversity, thereby providing an important fitness advantage (Eliezer et al., 2019). To determine if the inherited associative memory confers a fitness advantage to the progeny, we quantified survival rates of both trained P0 animals as well as of their F1 progeny. For this, we odor-evoked the stressful memory using the conditioned stimulus IAA, and allowed the stress response to develop for two hours, after which we subjected the worms to heat shock (37 °C) that typically kills ~50% of the worm population (**Figure 2A**).

**Figure 2.**
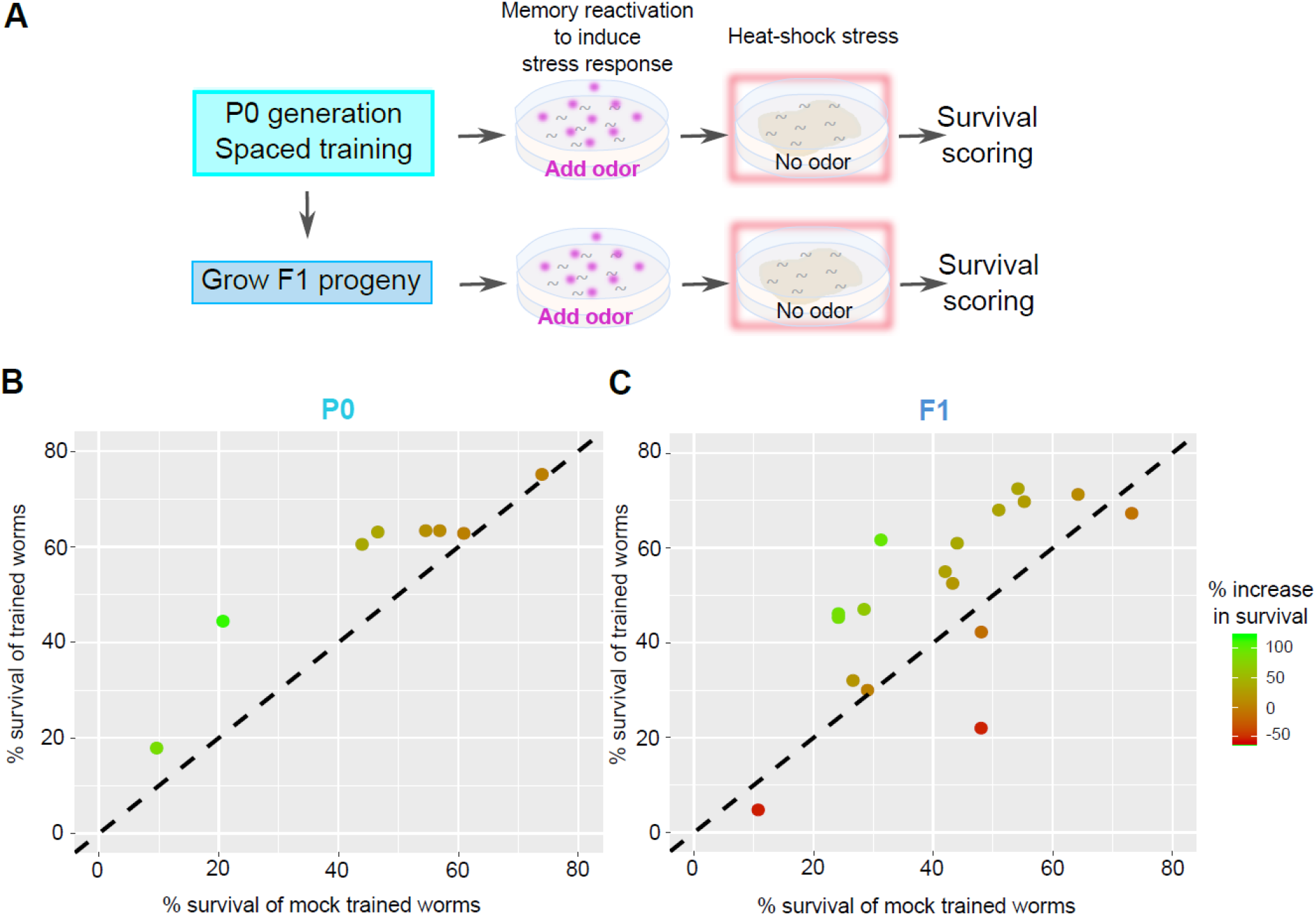
Reactivation of the inherited stressful memory increased survival chances of F1 descendants when facing stress. (**A**) The experimental design to quantify survival rates. P0-generation worms were trained (or mock-trained) to associate IAA with starvation. Their F1 progeny were grown under normal conditions (as in Figure 1A). One-day adult P0 and F1 worms were exposed to the conditioned stimulus IAA to reactivate the memory and induce the stress response. Following two hours, the worms were exposed to heat stress (37 °C), and scored for survival the following day. (**B-C**) Survival rates of trained P0 (B) and their F1 descendants (C), were significantly higher than their mock-trained counterparts when facing heat stress following memory reactivation. Note that only the P0 generation worms were trained. Each data point represents one experimental repeat in which ~50 trained and mock-trained animals were scored. The color of the data point indicates the % increase in survival, such that it is positive above the y=x dotted line. For P0 (B), N=8, p=0.0035 (paired t-test). For F1 (C), N=17, p=0.001 (paired t-test).

Survival chances of trained P0 worms were significantly higher than those of their mock-trained counterparts (**Figure 2B**). Furthermore, F1 progeny of trained P0 worms also showed significantly higher survival rates when compared to survival rates of descendants of the P0 mock-trained worms (**Figure 2C**). Thus, a mere reactivation of the inherited aversive memory allowed the F1 worms to prepare for imminent adversities in advance, and by this, to increase survival chances. These results demonstrate that parental associative memories can be inherited to endow the offspring with a valuable fitness advantage.

We next studied the molecular mechanisms by which these associative memories are transferred to the F1 progeny. When training P0 hermaphrodites (**Figures 1-2**), the memory information could had been transmitted via the sperm, the oocyte, or both. To elucidate which gametes carry the information, we trained either males or hermaphrodites, crossed them with naïve partners, and assayed the resulting F1 progeny for memory inheritance.

Interestingly, offspring of naive hermaphrodites that were crossed with trained males carried the capacity for memory reactivation (**Figure 3A**). In contradistinction, progeny of trained hermaphrodites crossed with naive males failed to show odor-evoked memory reactivation stress responses. Importantly, F1 descendants of a cross between trained males and trained hermaphrodites retained the capacity for odor-evoked memory reactivation, indicating that mating events do not impair memory inheritance (**Figure 3A**). Together, these results indicate that the memory information resides exclusively in the sperm of the trained parent.

**Figure 3:**
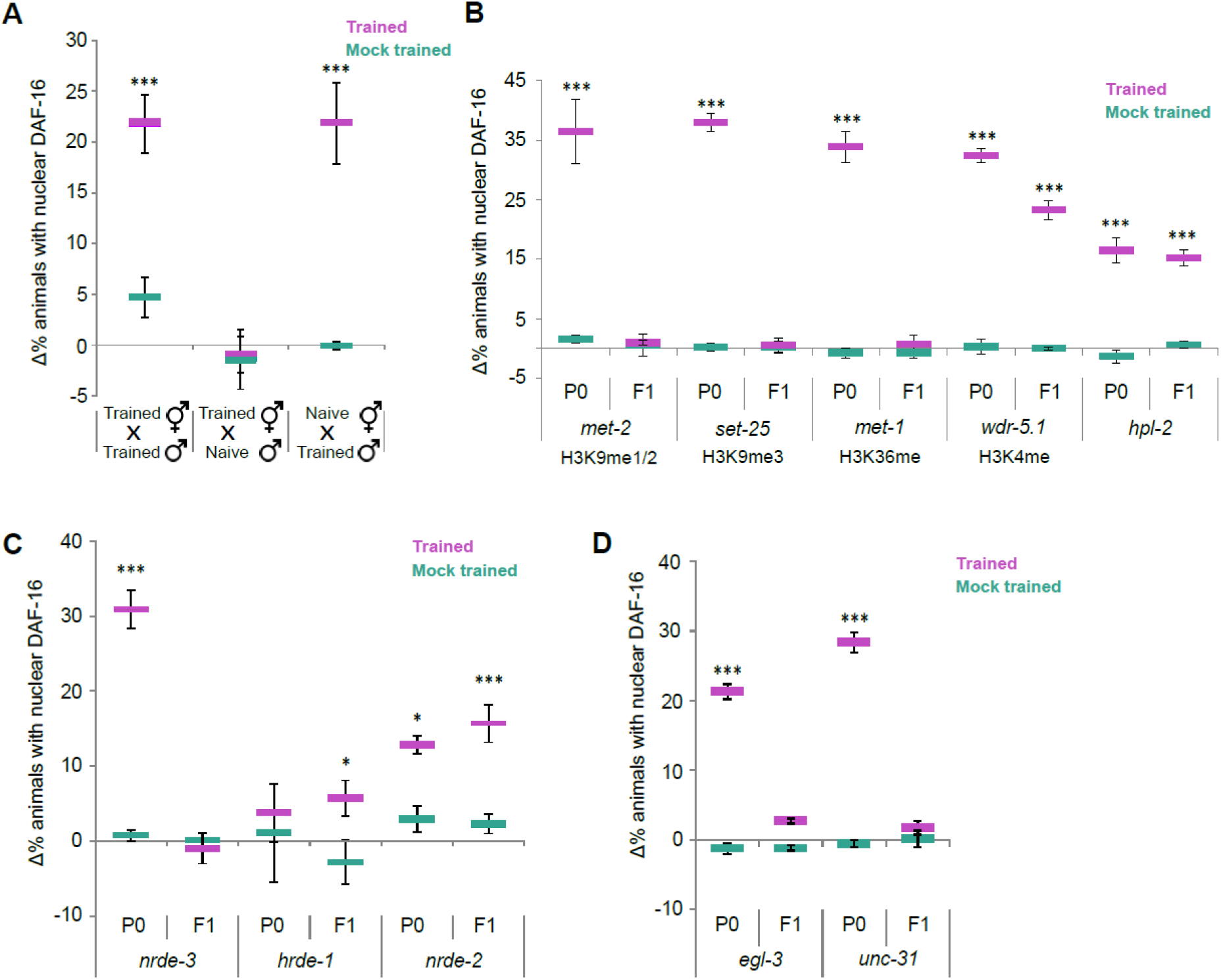
The associative memory is transmitted through the sperm, and depends on H3K9 and H3K36 methylation, the Argonaute NRDE-3, and neuropeptide secretion. (**A**) The memory is transmitted to progeny through the sperm only. Odor-evoked memory reactivation induced a rapid translocation of DAF-16/FOXO to cells’ nuclei only in F1s which were the progeny of either: (i) trained hermaphrodites crossed with trained males; (ii) trained males crossed with naïve hermaphrodites. F1 progeny of trained hermaphrodites crossed with naïve males lacked the associative memory. (**B**) H3K9 and H3K36 methylations are required for the inheritance of the associative memory. While trained P0 mutants, defective in H3K9 methylation (*met-2* and *set-25)* or H3K36 methylation (*met-1)* showed intact odor-evoked memory reactivation responses, this memory was not inherited by their F1 descendants. Mutants, defective in either *hpl-2* or *wdr-5.1,* showed reduced learning and memory abilities in both the trained P0 generation and their F1 progeny. (**C**) The Argonaute NRDE-3 is required for memory inheritance. While trained *nrde-3* P0 mutants formed and retrieved the memory, their F1 descendants failed to induce the stress response following memory reactivation. Mutants, defective in the Argonaute *hrde-1,* showed impaired memory formation and/or retrieval abilities in both the trained P0 and F1 generations, thus precluding inference of their role in the inheritance. *nrde-2* mutants showed reduced learning and memory abilities in both the trained P0 generation and their F1 progeny. (**D**) Neuropeptide signaling is required for memory inheritance. Trained P0 mutants of either *egl-3* or *unc-31* showed intact levels of memory retrieval, but this capacity was lost in their F1 progeny. In all panels, each data point is the mean±SE of 3–5 independent experimental repeats, each scoring ~50 animals. Significance comparisons are between trained and mock-trained animals. *p<0.05, ***p<0.0005 (proportion test).

In *C. elegans,* small RNAs and histone modifications constitute the main mechanisms underlying transgenerational epigenetic inheritance (Rankin, 2015; Rechavi and Lev, 2017; Minkina and Hunter, 2018; Weiser and Kim, 2019; Liberman et al., 2019; Perez and Lehner, 2019). We therefore analyzed the capacity to inherit associative memories in mutant animals, defective in key genes that function in these pathways. Since inheritance is mediated via either the male or the hermaphrodite sperm (**Figure 3A**), we simplified the following set of experiments by using hermaphrodites, leveraging on their self-fertilization reproduction.

We found that F1 mutants, defective in mono/di H3K9 methylation (*met-2)* and trimethylation (*set-25),* failed to inherit the memory, although the trained P0 mutant parents showed intact capacities to form and reactivate the memory (**Figure 3B**). Similarly, trained P0 mutants, defective in H3K36me (*met-1),* showed intact memory formation and retrieval abilities, but their mutant progeny failed to inherit the memory. In contrast, two other histone-related genes, namely, H3K4 methyl-transferase (WDR-5.1) and the heterochromatin protein, HPL-2, which binds methylated histones, do not seem to play a role in memory inheritance, as memory reactivation levels in both trained P0 mutants and their F1 descendants, are comparable (although lower compared to WT levels, **figure 3B**).

Next, we analyzed key genes in the small RNA pathways that were shown to function *C. elegans* transgenerational inheritance. F1 mutants defective in NRDE-3, an Argonaute responsible for transferring small RNAs from the cytoplasm to the nucleus in somatic cells (Guang et al., 2008), showed impaired memory inheritance, while the trained P0 generation showed WT levels of memory formation and retrieval (**Figure 3C**). In contrast, mutants defective in NRDE-2, the nuclear dsRNA-induced RNAi factor responsible for maintenance of transgenerational inheritance of small RNAs (Guang et al., 2008; Gu et al., 2012), showed intact capacities for memory retrieval in both trained P0 worms and their F1 descendants, suggesting that this gene is not involved in memory inheritance. Mutants in HRDE-1, a key nuclear Argonaute for transgenerational inheritance in *C. elegans* (Buckley et al., 2012), were defective in memory processes already at the trained parental generation, thus precluding from assessing the involvement of this gene in inheritance of associative memories.

We also speculated that neuronal secretion signals may be at play in the inheritance processes of associative memories. We therefore analyzed memory inheritance in two different mutants (*unc-31* and *egl-3)* that are defective in the general neuropeptide secretion pathways. Notably, while the parental trained generation showed intact WT levels of memory retrieval, F1 worms showed no memory retrieval capacities, suggesting that neuropeptides are also involved in the transmission of the associative memory (**Figure 3D**). These results place H3K9 and H3K36 methylation, the Argonaute NRDE-3, and neuropeptides as key factors that underlie inheritance of associative memories.

The associative memory paradigm used herein couples between starvation and the odorant IAA. IAA is sensed primarily by the bilateral AWC^ON^ and AWC^OFF^ chemosensory neurons (Chalasani et al., 2007; Zaslaver et al., 2015), which also participate in coding and storing the memory in the parental trained generation (Eliezer et al., 2019). We therefore asked whether these neurons also participate in storing the memory in the F1 generation?

To address this question, we used two transgenic lines, each exclusively expressing channelrhodopsin in either AWC^ON^ or AWC^OFF^. Remarkably, the mere light activation of AWC^OFF^ in the trained P0 worms as well as in their F1 descendants, sufficed to induce a rapid stress response (**Figure 4A**). In contrast, light-activation of AWC^ON^ led to a rapid activation in the trained P0 generation, but not in their F1 descendants. Moreover, inhibiting AWC^OFF^ activity using the inducible histamine-gated chloride channel during memory retrieval, abrogated memory retrieval in both trained P0 worms and their F1 offspring (**Figure 4B**). These results indicate that while both AWC neurons are part of the memory ensemble in the parental generation, in the offspring, only the AWC^OFF^ neuron participates in storing the memory and in inducing the systemic stress response.

**Figure 4.**
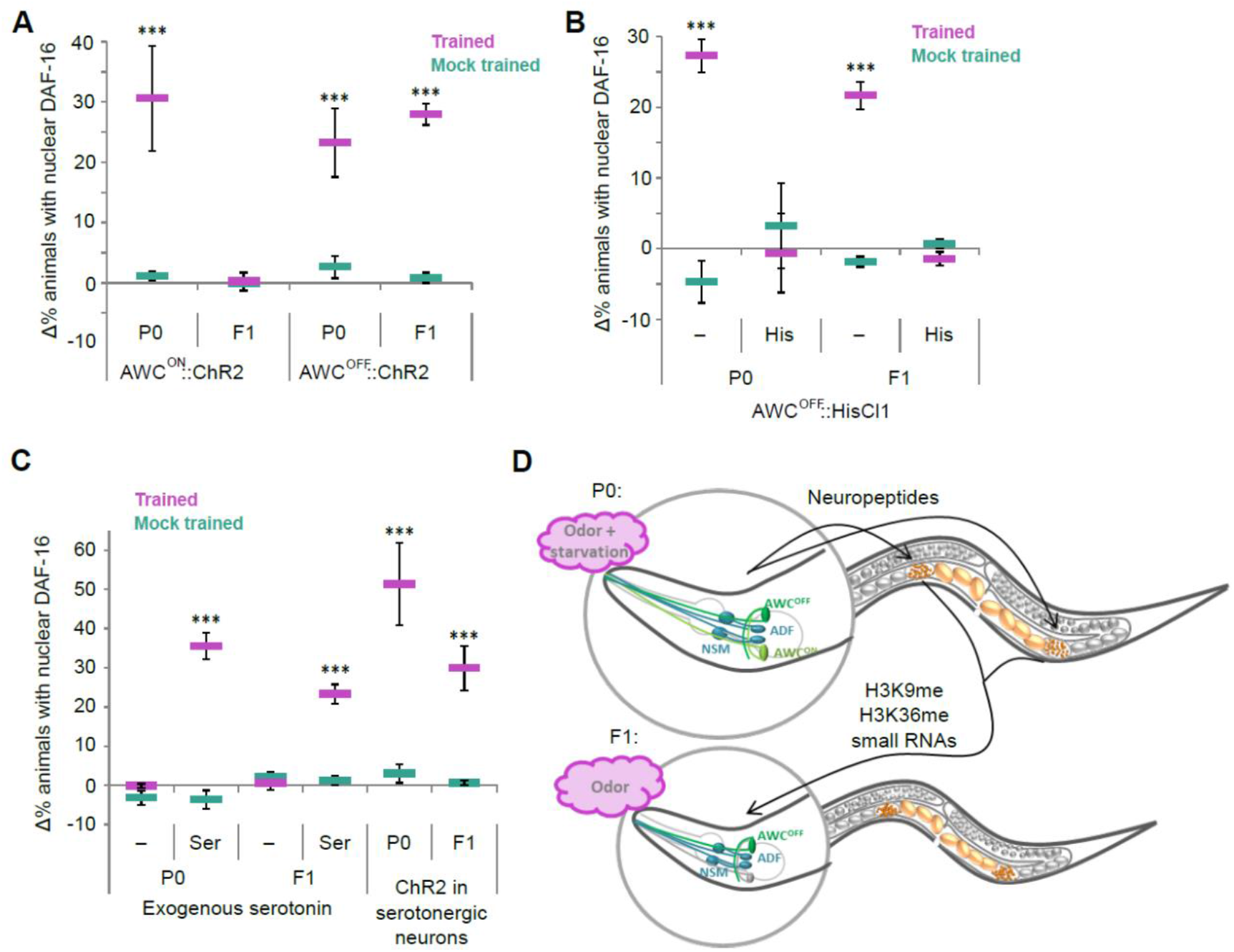
Partial overlap in the neurons that store the memory in the parental trained generation and their F1 descendants. In both, serotonin mediates the systemic stress response. **(A)** Light-activation of either the AWC^OFF^ or the AWC^ON^ neurons induced the stress response in the trained P0 generation. However, in the F1 generation, light-activation of the AWC^OFF^ neuron, but not of the AWC^ON^, induced the stress response. **(B)** The AWC^OFF^ neuron is required for memory reactivation and induction of the stress response in both the trained P0 generation and their F1 descendants. Exclusive inhibition of the AWC^OFF^ neuron during odor-evoked memory reactivation abolished induction of the stress response in both the parental trained generation and in their F1 progeny. **(C)** Serotonin mediates the systemic stress response following odor-evoked memory reactivation. Exposure to exogenous serotonin (without odor) induced DAF-16/FOXO nuclear translocation. Similarly, light-activation of the serotonergic neurons (ADF, NSM and HSN) induced a rapid systemic stress response in both the trained P0 worms and in their F1 descendants. In all panels, each data point is the mean±SE of 3–5 independent experimental repeats, each scoring ~50 animals. Significance comparisons are between trained and mock-trained animals. ***p< 0.0005 (proportion test). **(D)** A proposed model for inheritance of associative memory. In the trained P0 generation, the aversive memory can be induced by the exclusive activation of either the AWC^ON^ or AWC^OFF^ chemosensory neurons, suggesting that these neurons are part of the memory ensemble. Neuropeptides, small RNAs, and histone methylations (H3K9 and H3K36) work in concert to modulate the sperm state, and by this, transmit the associative memory information to the somatic cells of the next generation (e.g., the AWC^OFF^ and the serotonergic neurons). Notably, while essential for memory inheritance, these epigenetic factors are not required for memory formation in the parental trained generation.

In trained animals, serotonin acts downstream to the AWC neurons to mediate the rapid systemic stress response following memory reactivation (Eliezer et al., 2019). We therefore asked whether serotonin also mediates the systemic stress response in the progeny of the trained P0 animals. Indeed, applying external serotonin induced a rapid stress response in trained P0 animals as well as in their F1 offspring (**Figure 4C**). Moreover, light activation of the serotonergic neurons also induced a rapid stress response in both, the trained P0 worms and their F1s offspring (**Figure 4C**). Taken together, the same sensory neuron, AWC^OFF^, that acquired and stored the memory in the trained parental generation also stored the memory information in the progeny. Furthermore, once the memory was reactivated, serotonin signaling mediated the systemic stress response in both the trained parental generation and their F1 offspring.

## Discussion

Here we demonstrated that associative memories can be passed through generations. Analyses of mutant strains (**Figure 3**), together with functional interrogation of individual target neurons (**Figure 4A-C**), provide a possible mechanism for the inheritance (**Figure 4D**): The trained P0-generation animals form the associative memory where both AWC neurons participate in its storage. This memory is passed on to the next generation, but the capacity to induce the stress response is retained in the AWC^OFF^ neuron only.

How could the memory information pass from the parental somatic neurons to reside in the neurons of their progeny? A combination of small RNAs, histone modifications, and neuropeptides secretion mediate this process. We identified the Argonaute NRDE-3 as well as H3K9 and H3K36 methylations as key to the inheritance of associative memories (**Figure 3B-C**). As small RNAs operate in concert with chromatin modifiers (Gu et al., 2012; Rechavi and Lev, 2017; Weiser and Kim, 2019), it is possible that memory-specific endo-small RNAs direct the corresponding methyl transferases to establish loci-specific histone marks. In particular, NRDE-3 expression in developing embryos is essential for intergenerational inheritance (Guang et al., 2008), so it is plausible that RNAs are transferred to (or produced in) the sperm to establish epigenetic modifications in the developing embryos (Stoeckius et al., 2014b, 2014a).

While memory formation in the parental trained generation did not depend on intact neuropeptide secretion, the inheritance of the memory did (**Figure 3D**). Neuropeptides, and particularly the insulin-like peptides, are secreted based on the animal’s satiety conditions and often reflect its physiological state (Li, 2008). An intriguing possibility is that the inherited associative memory is established by integrating two seemingly unrelated cues: starvation that is signaled via selective neuropeptides, and the olfactory cue that is encoded by specific small RNAs targeting specific chromatin regions. Indeed, both neuropeptides and small RNAs were shown to be transferred from somatic tissues to the germline to affect the progeny (Devanapally et al., 2015; Burton et al., 2017; Posner et al., 2019; Perez et al., 2020; Kaletsky et al., 2020), and olfactory adaptation in the AWC neurons was shown to be mediated by NRDE-3 (Juang et al., 2013). Furthermore, stress-induced serotonin signaling modifies germ cells’ chromatin to promote their survival (Das et al., 2020).

In the parental trained generation, both AWC^ON^ and AWC^OFF^ neurons were potent to induce the stress response (Eliezer et al., 2019). In the progeny, however, only the AWC^OFF^ neuron retained the capacity to initiate the stress response. This indicates that the ensemble of memory-storing neurons only partially overlaps between generations. In fact, it is plausible that inherited memory includes only the set of neurons required for memory reactivation and excludes neurons that merely underwent experience-dependent changes in the trained parental generation. This partial overlap in neural ensembles may also explain the difference in the behavioral outputs, where the trained P0 generation withdrew from IAA while their F1 descendants did not (**Figure 1E**).

We consistently analyzed memory capacity in F1 animals that hatched from eggs laid 24 hours post the last training step (see Methods). As a fertilized egg is typically laid within three hours post fertilization (Hall et al., 2017), the analyzed F1 animals were likely to be in their germline pre-fertilized state during training. When worms were trained during the fourth larval stage (a single training step), robust associative memories were formed only in the P0 trained animals, but their progeny did not inherit the associative memory (**Supplementary figure 4**). This suggests that epigenetic marks take place in germ cells early on during gonad development.

Surprisingly, the associative aversive memory was also transmitted to the F2 generation (**Figure 3A**). F2s did not exist during the training period of the P0 generation, not even in their germ-cell state. They experienced the odorant IAA only once during their pre-fertilized state upon memory reactivation of their F1 parents (**Figure 1E**). This odor-evoked memory reactivation was performed on well-fed F1s that never experienced starvation (Methods), thus precluding the possibility of coupling between IAA and starvation. The transfer of the associative memory to the F2 generation suggests a trans-generational inheritance mechanism that does not depend on the direct exposure of the germline to the adverse conditions (Dias and Ressler, 2014; Dias et al., 2015; Perez and Lehner, 2019). An alternative intriguing possibility is that reactivation of the stressful memory is necessary for the transmittance, although F3-generation animals did not inherit the memory following memory reactivation of their F2 parents (**Figure 2E and supplementary figure 3**). In that respect, it may be interesting if inheritance of associative memories, like other transgenerational processes, is actively regulated to be limited to few generations only (Houri-Ze’evi et al., 2016).

Sperm, but not oocytes, transmitted the epigenetic memory (**Figure 3A**). In mammals, epigenetic inheritance was shown to be mediated by sperm, but the strong bias to perform experiments in male rodents precludes definite conclusions regarding oocytes involvement in such processes (Dias and Ressler, 2014). In *C. elegans* nematodes, exclusive inheritance via the sperm may have evolved due to their population dynamics constraints. *C. elegans* worms are androdioecious (Haag, 2005): Under non-stressful conditions, hermaphrodites make >99% of the population and the rest are males. However, upon stress, the relative fraction of males significantly elevates, presumably to increase probability of cross fertilization, and by this, genotypic diversity (Anderson et al., 2010). Thus, cross mating with males that developed through harsh conditions increases the likelihood that their adverse life history will be transmitted to subsequent generations. Moreover, the males’ sperm outnumbers and outperforms the hermaphroditic sperm, so cross fertilization becomes advantageous as it ensures a wider distribution of the adverse information among the descendants. Indeed, following mating, the sperm contributes to the 1-cell embryo a significant amount of mRNAs, particularly of endogenous siRNAs and piRNAs (Stoeckius et al., 2014b, 2014a). In the event of self-fertilization, the epigenetic memory can still be transmitted via the hermaphrodites’ own sperm (**Figure 3A**).

In summary, here we established that associative memories are inheritable and proposed a mechanism for this process. The inherited associative stressful memory encodes physiological protective instructions that endow the progeny a fitness advantage, thereby providing the evolutionary basis for the emergence of this valuable capacity in the animal kingdom.

## Materials and Methods

### Strains and growth conditions

A full list of the strains used in this study is available in supplementary Table 1. All strains were maintained and grown under standard conditions (Brenner, 1974), unless otherwise indicated, as for example during training or odor-evoked memory retrieval.

### Training the P0 generation to associate an odorant with starvation

We trained age-synchronized P0 generation worms by bleaching gravid hermaphrodites to extract fertile eggs (Stiernagle, 2006). The extracted eggs were seeded on NGM plates that were pre-coated with OP50 bacteria. Following ~18 hours in 20 °C, when animals reached the first larval stage (L1), we washed them off the plates with an M9 buffer. We then repeated the washing steps three more times to discard any bacterial residues and initiated the training procedure.

To form the association between the conditioned stimulus isoamyl alcohol (IAA) and starvation, we used a spaced-training paradigm that overall lasted one week. First, the washed L1 worms were transferred to bacteria-free NGM plates (starvation plates), either in the presence of the odorant IAA (trained animals), or in its absence (mock-trained animals). The starvation plates contained 2.5-3% agar to minimize worm burrowing, and ampicillin (100 μg/mL) to prevent possible contamination and bacterial growth. The IAA odorant was diluted 1:1000 and was added by applying 9 equally-distant 5 μL drops on the inside face of a 50 mm plate’s lid (the trained worms plate).

Following 24 hours, the worms were washed off the plates and transferred to OP50-seeded NGM plates for a period of ~6 hours to allow recovery from starvation. A second training was then imposed by washing the worms three times in M9 and transferring them (trained and mock-trained groups) to starvation plates for overnight conditioning in the presence or absence of IAA. At this stage, the worms experienced starvation as L2-L3 larvae. Following 18 hours of starvation, the worms were washed off the plates and allowed to recover on food (~6 hours), after which they were washed again and placed on starvation plates (with or without IAA) for an additional 18 hours (third starvation). We then recovered the worms for 4-8 hours on food. After this step, the majority of the worms reached the 4th larval stage (L4).

We then subjected these L4 worms to a fourth final ~65 hours long starvation (with or without IAA), followed by recovery on OP50-seeded NGM plates for 24 hours. By this time, the worms reached the young adult stage and started to lay eggs. About 60-70 worms, from each of the trained and the mock-trained adult P0 groups, were randomly selected to validate successful training and for further analyses.

### Retrieval of the F1 generation worms

We allowed the trained/mock-trained P0 generation to lay eggs for 24 hours during the last recovery period, after which we washed the plates with M9 to remove the P0 worms as well as F1 descendant larvae that already hatched. Following the washing step, we verified that only F1 eggs remained on the plate. Notably, washing off larvae that hatched during the 24 hours after the recovery period ensured that the F1 progeny remaining on the plate for analysis were still in the unfertilized germ state during the training of the P0 worms. After the eggs hatched, the F1 generation worms were collected and transferred to fresh OP50-seeded NGM plates for further growth. Once reaching adulthood, about 60-70 F1 generation worms from both trained- and mock-trained P0 worms were randomly selected for analysis.

### Retrieval of the F2- and F3-generation worms

For F2 generation analysis, we allowed the F1-generation worms to lay eggs, and then washed the worms off leaving on the plate unhatched eggs only. After the eggs hatched, the F2 generation worms were collected and transferred to fresh OP50-seeded NGM plates for further growth. Upon reaching adulthood, we randomly picked 60-70 of the F2 worms (descendants of each trained and mock-trained P0 worms) for further analysis (Figure 1C inset).

We also analyzed F2 (F3) generations that were descendants of F1 (F2) worms which were pre-selected for showing odor-evoked stress responses (Figure 1C and supplementary figure 1). For this, we challenged F1- and F2-generation worms with IAA for 20 minutes and transferred ~20 stressed worms to a fresh OP50-seeded plate. The worms were left for ~6 hours on the fresh plate in order to lay eggs, after which they were removed leaving laid eggs only. This procedure was done for F1 and F2 descendants of both trained and mock-trained P0 worms (again, only the P0 worms were trained). Subsequent F2 and F3 generations were grown to adulthood and then analyzed.

### Mating procedures of trained and naive P0 worms

To determine whether the inheritance is paternal or maternal, we used the ZAS394 strain, which, along with the DAF-16 translational reporter (*daf-16*::DAF-16::GFP (Henderson and Johnson, 2001)), also expresses the pan-neuronal red fluorescent marker (*rab-3*::tagRFP-NLS (Fry et al., 2016)) in the background of *him-5(e1490)* to increase male counts in the population. We trained either TJ356 (for maternal inheritance) or ZAS394 (for paternal inheritance), and after 24 hours of recovery period, we transferred 50 ZAS394 males (either naive or trained), and 15 TJ356 hermaphrodites (either trained or naive, respectively), to a large mating plate (at least 5 large mating plates per experiment). We allowed the worms to mate, and after three days we picked *rab-3*::tagRFP-NLS expressing offspring, to verify that they are the result of a successful mate. Those descendants were analyzed upon reaching adulthood as the F1 generation.

### Quantifying memory-induced stress responses based on FOXO/DAF-16 translocation to cells’ nuclei

Here, we followed the detailed procedures provided in (Eliezer et al., 2019). To quantify the induction of the stress response, we used worms expressing a translational fusion of the general stress response transcription factor FOXO/DAF-16 (*pdaf*-16::DAF-16::GFP (Henderson and Johnson, 2001)). A worm was scored as stressed if we clearly identified at least six cells with DAF-16::GFP nuclear localization. The cutoff of six cells was carefully selected based on our previous extensive analyses showing that this is a conservative estimation of the stressed worms in the population (Eliezer et al. 2019). We focused on scoring DAF-16::GFP nuclear localization in the gonad sheath cells as this tissue was the first to show this spatial dynamic pattern.

Each experiment always included four groups of worms: trained and mock-trained groups, where each group was further sub-divided into a challenged group (in which the memory was reactivated using an odorant or optogenetically), and an unchallenged group. The non-challenged groups served as a measure for the baseline stress levels in the population, and these levels were subtracted from the stress levels quantified in the challenged groups. When worms did not initiate stress responses following memory reactivation, the % of stressed worms was minimal (1-3%), very close to the control non-challenged group of worms. This is why we obtained negative values when calculating the % of stressed mock-trained animals, or when assaying conditions/mutants that did not elicit stress in trained animals.

Importantly, following the last recovery step from starvation and prior to challenging the worms for memory reactivation, we verified that worms had fully recovered from the starvation-induced stress. This was done by visual inspection of the DAF-16::GFP protein, ensuring that its spatial localization within the cells was cytoplasmic (rather than nuclear).

All scoring procedures were done in a blind manner such that the examiner did not know the treatment that the worms underwent (e.g. trained or mock trained). Each scoring plate consisted of at least 50 animals that were inspected under a fluorescent binocular (MVX10, Olympus) with a high-zoom magnification (X300). Typically, each data point is the average of at least three independent experimental repeats performed on different dates.

For memory reactivation using the light-activated ChR2, we recovered the worms from starvation on plates pre-seeded with OP50 containing All-Trans Retinal (100 μg/ml, Sigma). Following the recovery period, we exposed the worms (while on the same recovery plates) to 488 nm blue light for 20 minutes before scoring. The light source was the LED coupled to the fluorescent binocular. Notably, the blue light on its own did not induce stress, as no nuclear DAF-16::GFP was observed in the mock-trained animals, nor in their F1 and F2 progeny.

To inhibit neural activity via the histamine-gated chloride channel (Pokala et al., 2014), we placed the worms on plates pre-seeded with OP50 supplemented with 10 mM histamine (Sigma). The worms were allowed to absorb the histamine for at least 40 minutes before applying IAA for memory reactivation.

In serotonin feeding experiments, the worms were placed on plates pre-seeded with OP50 mixed with 10 mM serotonin (Sigma). The worms were first inspected to verify that the mere handling and transfer did not cause DAF-16 to translocate to cells’ nuclei. The animals were then allowed to absorb the serotonin for 20 minutes, after which we scored for DAF-16 nuclear localization.

#### Avoidance assays

For behavioral assays, we used WT N2 worms. We trained the P0-generation to form an associative aversive memory as described above (Figure 1A), and individual trained or mock-trained animals (either of the P0 or the F1 generations) were transferred to unseeded plates, and allowed to adjust for ~2 minutes before the assay. First, we inspected when a worm performed a long forward run, and then presented it with IAA (1:1000 dilution in water) by spreading a thin stripe of the odorant, perpendicular and in front of its forward trajectory. Worms that stopped in front of the spread stripe and backed within three seconds were scored as ‘avoiding’; worms that only briefly stopped, or continued to crawl forward crossing the stripe were scored as ‘not avoiding’ (Figure 1F).

#### Fitness (survival) analysis

We trained WT N2 hermaphroditic worms, which constituted the P0 generation. To evoke the memory and induce the stress response prior to the heat shock, P0 or F1 worms were transferred to fresh plates without food and presented with IAA (10^-3^) for two hours by applying 9 equally-distant 5 μL drops on the inside face of the plate cover. The worms were then transferred to OP50-containing plates and subjected to a 37 °C heat shock, a stress that typically led to ~50% lethality. The P0-generation worms were heat shocked for 4.5-5.5 hours while the F1-generation worms were heat shocked for 3.5-4.5 hours. The extended heat shock period used for the P0 generation was due to their higher resistance to heat, presumably due to the intense stress (repeated starvations) that they underwent throughout their developmental stages. Since heat shock induces quiescence, it is difficult to assess mortality immediately after the heat shock. We therefore scored viability on the following day, when surviving animals were clearly motile, pumping, and withdrawing following a gentle touch on their head. Animals not showing any of these features were scored as dead.

#### Statistical analysis

We used the proportion test to infer statistical significance between the different groups and conditions. For this, we considered two pairs of proportions 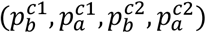, where each pair represents the proportion of the stressed worms before (*b)* and after (*a*) the challenge in the two compared conditions (*c1, c2; e.g.,* trained and mock trained). This analysis tests whether the difference between the first condition proportions 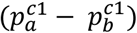 is likely to be sampled from the expected distribution of the difference between the second condition proportions 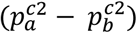.

Formally, the null hypothesis is defined as:

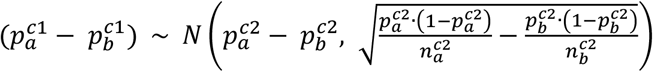

Where n is the total number of analyzed animals in this condition.

## Acknowledgements

We thank Itamar Harel and Yonatan Tzur for insightful comments on the manuscript draft. We are also thankful to Meital Oren, the Mitani lab, and the CGC for strains. The CGC is funded by the NIH Office of Research Infrastructure Programs (P40 OD010440). Our lab is supported by ERC (336803), ICORE (1200/12), and ISF (1300/17). AZ thanks Joseph H. and Belle R. Braun senior lecture chair and Greenfield chair in Neurobiology.

## Author contributions

ND, YE, LH, and SBE performed the experiments. EI developed statistical methods. ND, YE and AZ analyzed and interpreted the data. AZ, ND, and YE wrote the paper.

## Supplementary Figures and Tables

**Supplementary figure 1.**
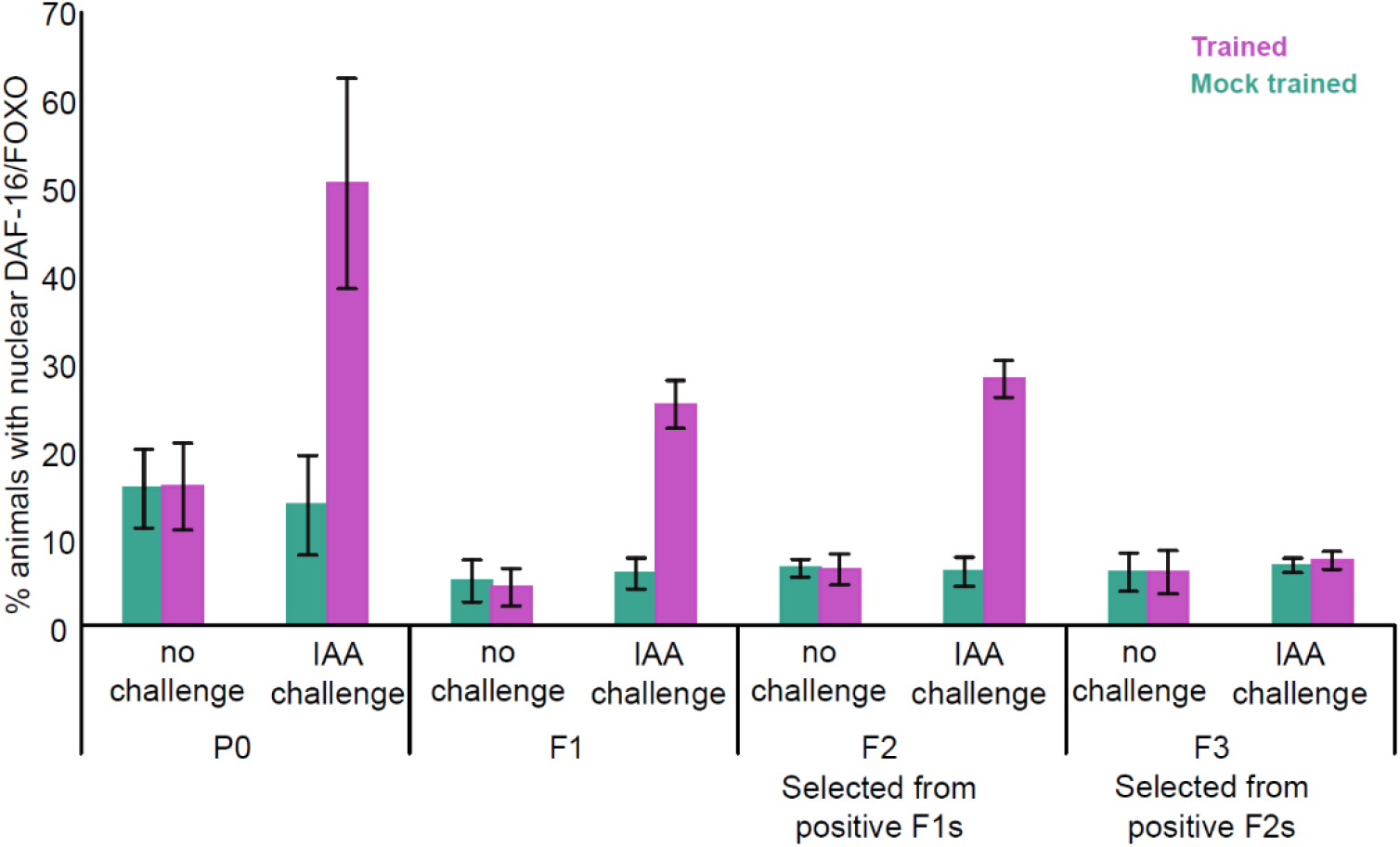
Aversive associative memories are trans-generationally transferred to the F1 and F2, but not to the F3, generations. Odor-evoked memory reactivation induced the translocation of DAF-16/FOXO to cells’ nuclei in the F1 and F2 generations, but not in the F3 generation. Animals were scored as having nuclear DAF-16/FOXO if at least six cells (primarily gonad sheath cells) were detected with nuclear DAF-16/FOXO within 20 minutes following memory reactivation. This cutoff is a conservative estimation and is based on our previous rigorous analysis (Eliezer et al., 2019). For each generation, we scored trained and mock-trained animal groups, where each group was scored before and after the IAA challenge. The non-challenged groups served as control to assess the overall stress in the population without evoking the stressful memory for both trained and mock-trained animals. These data provide the raw values on which figure 1 was compiled: For each group we subtracted the % of worms with nuclear DAF-16/FOXO before the challenge with IAA from the % of worms with nuclear DAF-16/FOXO after the challenge with IAA. The same subtracted values were used in all other figures shown in the main text, and this is also the reason why negligible % of stressed worms sometimes results in negative values. The animals scored for F2 and F3 generations were progeny selected from F1 and F2-generation animals that showed stress induction following odor evoked memory retrieval. F1 animals were not preselected and were randomly collected from all P0-trained animals. Each bar is the average of at least three independent experimental repeats, each scoring ~50 animals. Error bars indicate SEM.

**Supplementary figure 2.**
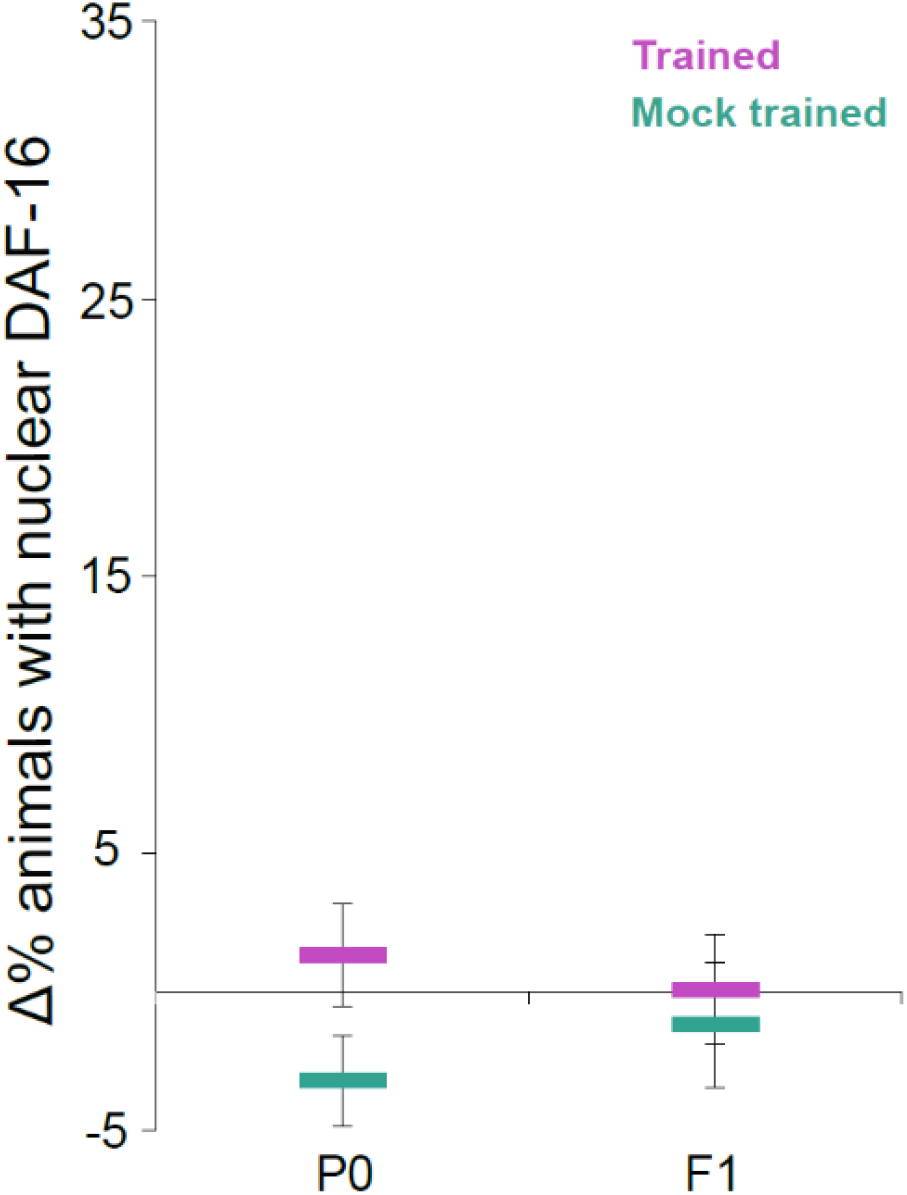
Worms trained to associate IAA with food do not initiate stress responses upon exposure to IAA. P0-generation worms were spaced trained to associate IAA with food. The F1 generation were grown in normal conditions on food and were never exposed to IAA. Odor-evoked memory reactivation did not induce a stress response in the trained P0 generation, nor in their F1 progeny, as we did not observe DAF-16/FOXO translocation to cells’ nuclei. Each data point is the mean of three independent experimental repeats, each scoring ~50 animals. Error bars indicate SEM.

**Supplementary figure 3.**
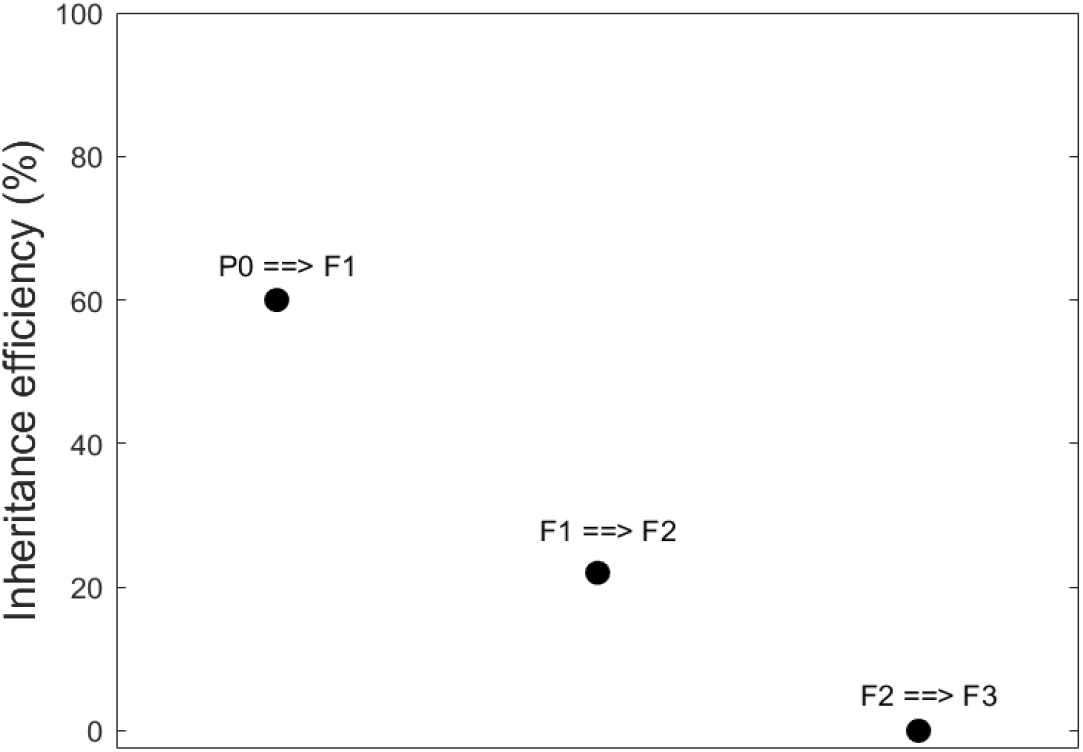
The efficiency of memory inheritance decreases through generations. The analysis herein relies on the data provided in figure 1D. ~33% of the trained P0-generation animals acquired the memory (or at least had shown the capacity for memory reactivation, figure 1D). In the next F1-generation, ~20% of the animals exhibited memory reactivation abilities. Since only ~33% of their parents acquired the memory from the first place, it is estimated that inheritance efficiency from the trained-P0 generation to their F1 descendants is ~20/33% (~60%). With this inheritance efficiency, we would expect that 60% of the 20% F1s that inherited the memory will also acquire the memory (thus, 12%). However, we find that the % of F2 animals that showed memory reactivation abilities was close to zero, suggesting that the inheritance efficiency from the F1 generation to their F2 descendants is lower than 60% (Figure 1D, inset). When analyzing F2 animals that were derived exclusively from memory-positive F1 animals, 22% of the F2 animals showed memory reactivation capacities. This indicates a ~22% efficiency in inheritance from the F1 to the F2 generation. Similarly, when analyzing F3 animals that were derived from memory-positive F2 animals only, we would expect to find that 22% of the F3 generation acquired the memory. However, once again, the % inheritance dropped to undetectable levels, presumably to 0%, suggesting that the memory inheritance lasts for two generations only.

**Supplementary figure 4.**
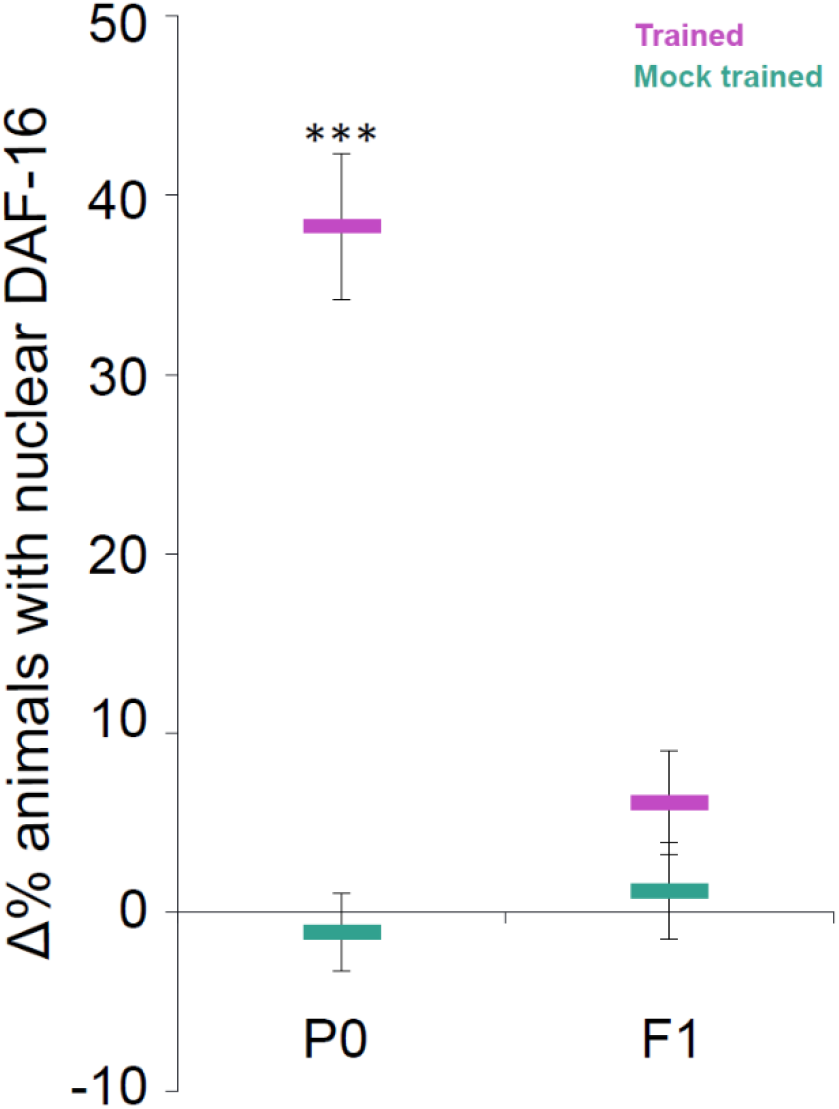
A single training paradigm forms robust memories in the P0 trained generation, but this memory is not transferred to their progeny. P0 generation hermaphrodites were trained to associate the odorant IAA with starvation during their fourth larval stage. Odor-evoked memory reactivation of trained animals induced a rapid stress response, as indicated by the translocation of DAF-16/FOXO to cells’ nuclei. However, this associative memory did not transfer to the F1 generation. Each data point is the mean of three independent experimental repeats, each scoring ~50 animals. Error bars indicate SEM. ***p < 0.0005 (proportion test).

**Supplementary Table 1.** Reagents and resources used in this study.

| REAGENT or RESOURCE | SOURCE | IDENTIFIER |
| --- | --- | --- |
| **Chemicals** |  |  |
| Histamine dihydrochloride | Sigma-Aldrich | CAT#H7250 |
| Serotonin creatinine sulfate monohydrate | Sigma-Aldrich | CAT#H7752 |
| Isoamyl alcohol | Sigma-Aldrich | CAT# W205702 |
| Streptomycin | Sigma-Aldrich | CAT# S6501 |
| Ampicillin | Sigma-Aldrich | CAT#A9518 |
| **Bacterial Strains** |  |  |
| *E. coli*: OP50 | Caenorhabditis<br>Genetics Center | RRID: WB-STRAIN: OP50 |
| *E. coli*: OP50-1; Streptomycin resistant | Caenorhabditis<br>Genetics Center | RRID: WB-STRAIN: OP50-1 |
| ***C. elegans* strains** |  |  |
| *C. elegans* wild type | Caenorhabditis<br>Genetics Center | RRID: WB-STRAIN: N2 |
| TJ356: *zls356* [*daf-16p*::DAF-16a/b::GFP + *rol-6(su1006)*] | (Henderson and<br>Johnson, 2001) | RRID: WB-STRAIN: TJ356 |
| *him-5* (e1490) x TJ356 | (Eliezer et al., 2019) | ZAS204 |
| ZAS53 (*azrEx53*[*str-2*::ChR2-mcherry], *lite-1*(ce314)) x TJ356 | (Eliezer et al., 2019) | ZAS170 |
| ZAS105(*azrEx105[srsx-3::Chr2-cherry+PHA-1]*; *pha-1(e2123ts)*; *lite-1(ce314)*) x TJ356 | (Eliezer et al., 2019) | ZAS234 |
| ZAS253 (*azrEx253[srsx-3::HisCl1::SL2::NLS-mCherry + PHA-1]*, *pha-1(e2123ts)*, *him-5(e1490)*) x TJ356 | (Eliezer et al., 2019) | ZAS270 |
| *zls356[daf-16p::DAF-16a/b::GFP + rol-6(su1006)]* x *unc-31(e928)*; | (Eliezer et al., 2019) | ZAS178 |
| INV60006 (*lite-1(ce314)*; Ex[*ptph-1::Chr2-mCherry punc-122::GFP*]) x TJ356 | (Iwanir et al., 2016) | ZAS309 |
| ZAS204 x OH10690 (*otls356[rab-3p(prom1)::2xNLS::TagRFP]*) (Fry et al., 2016) | This study | ZAS394 |
| *set-25(n5021)* x TJ356 | This study | ZAS313 |
| *met-1(n4337)* x TJ356 | This study | ZAS316 |
| *met-2(n4256)* x TJ356 | This study | ZAS318 |
| *hpl-2(tm1489)\** x TJ356 | This study | ZAS284 |
| *wdr-5.1(ok1417)* x TJ356 | This study | ZAS328 |
| *nrde-2(gg91)* x TJ356 | This study | ZAS420 |
| *hrde-1(tm1200)\** x TJ356 | This study | ZAS339 |
| *nrde-3(tm1116)\** x TJ356 | This study | ZAS340 |
| *egl-3(n150)* x TJ356 | This study | ZAS182 |

| Software |  |  |
| --- | --- | --- |
| R Project for Statistical Computing | The R Foundation | RRID:SCR_001905 |
| RStudio Version 1.1.419 | RStudio Team<br>(2015) | RRID:SCR_000432 |
| ImageJ | Fiji | RRID:SCR_003070 |

\*These strains were provided by the Mitani lab through the National Bio-Resource Project of the MEXT, Japan.

